# In Silico Analysis of Effect of Non-synonymous SNPs associated with Permanent Neonatal Diabetes Mellitus on Stability and Structure of Human Insulin

**DOI:** 10.1101/2020.09.01.278002

**Authors:** Nishat Tabassum

## Abstract

**Motivation:** Proteins are the building-blocks of life. However with the deluge of data on protein sequences and their association with diseases, it is imperative to use computational tools to aid experimental analysis. Determination of the impact of disease-causing SNPs on protein’s structure is crucial in discovering how a disease affects the functions on a fundamental level and thereafter, determining potential drug targets.

**Results:** SNPs associated with the genetic disease permanent neonatal diabetus mellitus (PNDM) were studied to determine their impact on human insulin. Out of 16 missense variants, eight were predicted to be deleterious. 6 of the SNPs resulted in high structural differences (RMSD > 0.9, H bonds > 35). Stability changes were also determined.

## 1 Introduction

Single-nucelotide polymorphisms (SNPs) are the most frequent genetic variations. The human genome is known to contain 4-5 million SNPs (Shamir and Pevzner, 2011). Among them, non-synonymous SNPs bring about changes in the amino acid residues, which can have significant effect on the functional characteristics of the derivative proteins. They can therefore be associated with diseases. The stability of the protein, its active sites, interactions with other proteins are all dependent with the structure (Yates and Sternberg, 2013). Consequently, it is reasonable to assume that non-synonymous SNPs are likely to alter the structure of the native protein. Several studies have focused on determining the impact of deleterious SNPs on the structure of native proteins (Elkhattabi *et al*., 2019, Younus *et al*., 2018). Usually SNPs that arise in highly conserved positions can significantly affect the functional and structural characteristics of a protein, which makes these SNPs strong candidates for being associated with diseases (Younus *et al*., 2018).

Elkhattabi *et al*. (2019) studied the effect of SNPs on the structure of human resistin; out of 78 SNPs studied, 15 were predicted to be deleterious. 9 of these were found to exist in highly conserved regions, presenting significant impact on the structure.

A similar scheme was adopted to study the impact of deleterious SNPs on human insulin protein. Permanent neonatal diabetus mellitus (PNDM) is a genetic disease that affects infants in the first 12 months of their lives, necessitating continuous insulin treatment. Developmental delay, risk of diabetus mellitus in adulthood, hypoglycemia are acute complications that may arise if the disease is improperly managed (De León and Stanley, 1993). Computational analysis of deleterious SNPs not only supports the effort of finding true disease associations, but can be used to determine potential binding sites for a more targeted approach to treatment (Zhang, nd).

## 2 Methods

The methodology closely followed the original workflow discussed in Elkhattabi *et al*. (2019).

### 2.1 Dataset collection

The amino acid sequence of the insulin protein (Accession: P01308) was obtained from UniProt reviewed database (https://www.uniprot.org/uniprot). The entry has a curated list of the non-synonymous changes in the amino acid sequence that have been linked to permanent neonatal diabetus mellitus (PNDM) - a rare form of the disease that permanently affects children. From the dbSNP database, the SNPs responsible for the variants predicted to be responsible for the genetic disease were retrieved (https://www.ncbi.nlm.nih.gov/SNP). Table 1 shows the missense variants and their SNP IDs.

**Table 1.** Missense variants associated with PNMD and the corresponding SNPs

| Position | Variant | SNP ID |
| --- | --- | --- |
| 24 | A to D | rs80356663 |
| 29 | H to D | rs121908272 |
| 32 | G to R | rs80356664 |
| 32 | G to S | rs80356664 |
| 35 | L to P | rs121908273 |
| 43 | C to G | rs80356666 |
| 47 | G to V | rs80356667 |
| 48 | F to C | rs80356668 |
| 84 | G to R | rs8121908274 |
| 89 | R to C | rs80356669 |
| 90 | G to C | rs80356670 |
| 96 | C to S | rs80356671 |
| 96 | C to Y | rs80356671 |
| 101 | S to C | rs121908276 |
| 103 | Y to C | rs121908277 |
| 108 | Y to C | rs80356672 |

### 2.2 Determination of deleterious SNPs

A consensus web-based classifier named PredictSNP2.0 (Bendl *et al*., 2014) was used to predict whether the single nucleotide variants were in actuality associated with any deleterious effects. PredictSNP2.0 utilizes nine prediction tools to develop a consensus: SIFT, PolyPhen-1, PolyPhen-2, MAPP, PhD-SNP, SNAP, PANTHER, PredictSNP, and nsSNPAnalyzer.

### 2.3 Conservation of sequence

ConSurf web server (http://consurf.tau.ac.il/) was used to determine the conservation of the amino acid sequence. The web-based tool assesses the functional regions of the protein on the basis of conserved blocks that emerge in a multiple sequence alignment. Scores between 1 and 9 are assigned to each residue, with a score of 9 given to the most conserved position and 1 to the least.

### 2.4 Prediction of stability changes

MUpro tool was used to predict change in protein stability due to the non-synonynous SNPs (http://mupro.proteomics.ics.uci.edu/). The tool utilizes support vector machine and neural networks to obtain two predictions (Cheng *et al*., 2006); the accuracy is reported to be above 84%. The confidence score is from -1 to 1; <0 indicates decrease in stability, whereas >0 suggests that the protein stability increases due to the mutation.

### 2.5 Analysis and comparison of structure

The native structure was chosen as the nuclear magnetic resonance verified structure of highly stable human insulin (PDB ID: 5wbt.1.A) and mutant structures were modelled using the Swiss model platform (https://swissmodel.expasy.org). For each structure, it was ensured that the template chosen had >80% sequence identity among the templates and resulted in a model with a favorable QMEAN score. The QMEAN score is an estimate of the “degree of nativeness” of the structural features observed in the model. It indicates whether the QMEAN score of the model is comparable to what one would expect from experimental structures of similar size. QMEAN around zero indicates good agreement between the model structure and experimental structures of similar size. Scores of -4.0 or below indicate models with low quality (Waterhouse *et al*., 2018). UCSF Chimera (Pettersen *et al*., 2004) was then used to align the native structure with each of the mutant proteins in turn to check the RMS deviation and changes in hydrogen bonds. UCSF Chimera offers a platform for visualization and analysis of molecular structures.

## 3 Results

### 3.1 Prediction of deleterious SNPs

The protein entry in UniProt listed a total of 16 missense variants associated with PNDM. The SNP data for these 16 variants was retrieved in September 2019 from the NCBI SNP database. Only 8 were predicted to be deleterious by the PredictSNP2.0 consensus classifier (Table 2). The SNPs that were predicted to be non-deleterious by consensus were discarded from further analysis.

**Table 2.** Results of SNP analysis by PredictSNP

| dbSNP ID | PredictSNP2 | CADD | DANN | FATHMM | FunSeq2 | GWAVA |
| --- | --- | --- | --- | --- | --- | --- |
| rs80356664 | -87% | -52% | -72% | -56% | +62% | - |
| rs121908273 | -87% | -52% | -68% | -63% | +62% | - |
| rs80356667 | -87% | +74% | -60% | -83% | -62% | +52% |
| rs80356668 | -82% | +59% | +60% | -68% | +62% | -50% |
| rs80356669 | -82% | +76% | -70% | +63% | -62% | +52% |
| rs80356670 | -82% | -52% | -70% | +63% | -62% | +53% |
| rs80356671 | -77% | -60% | -75% | -69% | +60% | -55% |
| rs80356672 | -77% | +54% | -64% | -69% | -67% | +65% |

- denotes deleterious effect, + denotes beneficial effect

### 3.2 Analysis of conservation

ConSurf analysis revealed that the eight SNPs were located in regions under high conservation (Table 3). This is a strong indication of the impact of these SNPs on the structure and stability of the native protein.

**Table 3.** Conservation analysis by ConSurf

| Residue position | Conservation score | B/E |
| --- | --- | --- |
| G32 | 9 | E |
| L35 | 9 | B |
| G47 | 9 | B |
| F48 | 9 | B |
| R89 | 9 | E |
| G90 | 9 | E |
| C96 | 9 | B |
| Y108 | 9 | E |

B-Buried, E-Exposed

### 3.3 Effect of deleterious SNPs on stability

Using MUpro tool, the change in stability of the protein structure due to the eight missense SNPs were predicted. Results are shown in Table 4.

**Table 4.**
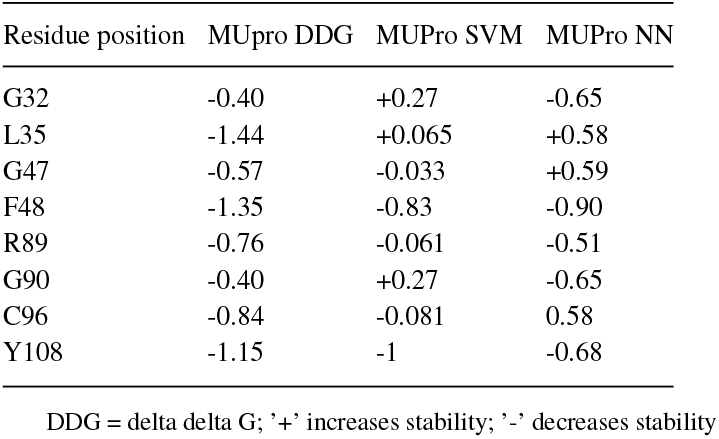
Stability prediction for mutations

### 3.4 Structural analysis

Swiss model web server was used to model the mutant structures; the model with the best QMEAN score, for which the template had more than 80% sequence identity were chosen. Table 5 lists the QMEAN scores and sequence identity of the templates chosen to model the proteins. Figure 1 shows the structure of the native protein. Using UCSF Chimera, each mutant model was superimposed on the native structure in turn to determine the root mean square deviation (RMSD) values, which measures the quality of the structural alignment. Higher the difference, greater the differences in the two structures and correspondingly higher probability of activity alteration (Elkhattabi *et al*., 2019). The total H-bonds were also compared between the structures, a difference indicating changes in folding and stability. The native protein had 35 H-bonds in its structure. Table 6 gives the RMSD values and total number of H-bonds in the mutant structures. Figure 2 shows the superimposition of the two mutant proteins giving the highest RMSD values. High variability in the loop regions can be noted.

**Table 5.** Swiss model template identity

| Protein | QMEAN score | Sequence identity (%) |
| --- | --- | --- |
| Native | -3.17 | 87.72 |
| G32 | -3.01 | 89.47 |
| L35 | -2.71 | 91.67 |
| G47 | -3.83 | 89.47 |
| F48 | -2.84 | 85.96 |
| R89 | -2.56 | 85.96 |
| G90 | -1.93 | 89.47 |
| C96 | -3.11 | 89.47 |
| Y108 | -2.60 | 89.47 |

**Table 6.** Results of superposition

| SNP | RMSD (Å) | No. of H-bonds |
| --- | --- | --- |
| G32 | 0.978 | 50 |
| L35 | 0.984 | 50 |
| G47 | 0.981 | 42 |
| F48 | 0.222 | 35 |
| R89 | 0.132 | 30 |
| G90 | 0.946 | 42 |
| C96 | 0.969 | 46 |
| Y108 | 0.944 | 38 |

**Fig. 1:**
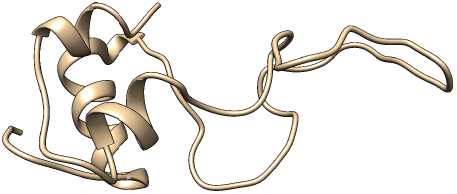
Model of Chain A of human insulin generated via Swiss-modelling.

**Fig. 2:**
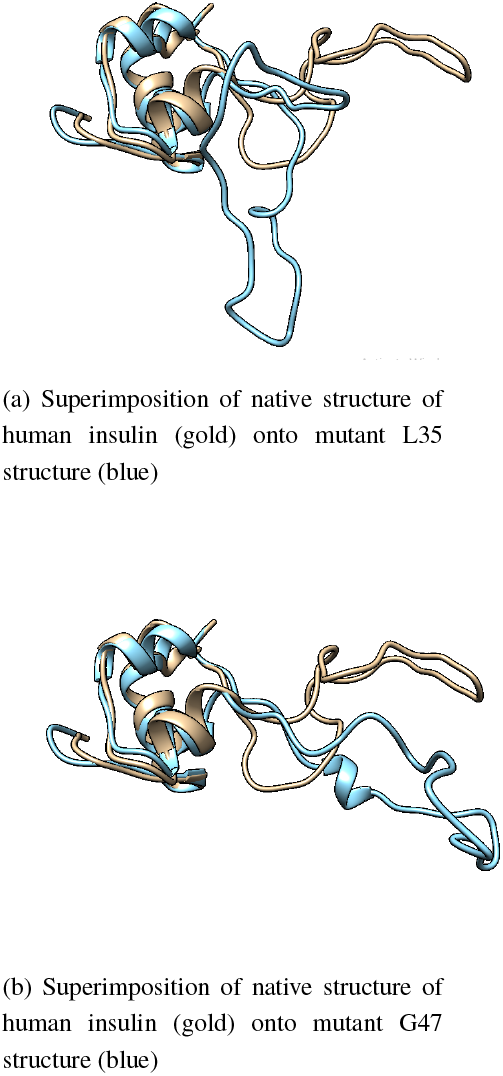
Structural differences between native and mutant proteins.

## 4 Discussion

Out of the 16 missense variants associated with PNDM, eight were predicted to be deleterious. This proves that computational tools can drastically reduce the scale of the problem. The analysis reveals that SNPs affect protein stability and structural aspects crucial to the protein’s integrity, such as hydrogen bonds. More sophisticated analyses would reveal changes in binding sites that render the protein inactive. From the eight deleterious SNPs, except F48 and R89, all other mutations resulted in high RMSD values (above 0.9), and these mutant structures had higher number of H bonds as well (above the number the native protein has). The mutations with RMSD values above 0.9 were predicted to have strong destabilizing effect on the protein. The high values of RMSD indicate significant differences in structure. H bonds determine how the protein folds, which affects the manner in which it interacts with other proteins and molecules (Claverie and Notredame, 2007). These residue positions having missense variants exist in the highly conserved regions of the protein. This lends credence to the fact that any changes in their identity is likely to have drastic effects on the functions of the protein. Biological data deluge is what prompted the development of *in silico* methods of analysis. Instead of testing thousands of SNPs, isolating deleterious ones using these tools would allow a more directed approach to epidemiological studies. Residues with highly deleterious variants could be possible candidates as drug targets (Younus *et al*., 2018). This analysis could be extended further to determine changes in the ligand-binding capability of the protein to definitively determine how changes in the structure affects functional characteristics.

## 5 Conclusion

In this study, computational proteomics tools were used to analyze various aspects of proteins. The first part of the study was used to examine the structural effects of deleterious SNPs associated with the disease PNDM. Structural alignment revealed differences in structure; highest RMSD was observed with SNPs that resulted in the greatest change in number of H bonds.

